# A long tail of truth and beauty: A zigzag pattern of feather formation explains the symmetry, complexity, and beauty of the peacock’s tail

**DOI:** 10.1101/2020.01.03.893776

**Authors:** Rama S. Singh, Santosh Jagadeeshan

## Abstract

One hundred and fifty years after the publication of Darwin’s sexual selection theory, the problem of the peacock’s train remains unsolved. Darwin assumed that the peacock’s long train was maladaptive and was the indirect effect of selection by female mate choice based on the train’s beauty. While a relationship between the feathers’ elaborate features and mating success has been shown, the concept of eyespots as the basis of female choice remains controversial. We examined the anatomical plan underlying feather development using museum specimens and observed a zigzag pattern of feather follicles that determined both the number and the hexagonal arrangement of eyespots on the train as well as, strikingly, the individual eyespots’ color rings. While the zigzag pattern explains the symmetry, complexity, and structural beauty of the peacock’s train, it also precludes individual eyespot variation. The only available variation in eyespot number is expected to be due to annual addition of new rows of 10/11 feathers as a function of age, giving rise to an inherently determined eyespot number. New insights show that eyespot number and feather length are developmentally correlated and an asymptotic function of a male’s age, that their effects on female choice would be confounded and inseparable, and that male vigor would be a crucial factor affecting male fitness. Females may not always choose males with the largest number of eyespots, as older males may lack vigor. We propose a multimodal model of female choice based on male size, vigor, and beauty where females see eyespot and train size not as separate traits but as one complex trait combining both. The new model may be able to explain conflicting results and why eyespot number alone may not be sufficient to explain female choice.

*Beauty is truth, truth beauty*.

Keats

## 1. Introduction

The Indian blue peacock’s, *Pavo cristatus*, elaborate and long tail (we use *tail* and *train* interchangeably, as appropriate-former to signify the long horizontal tail, the latter in expanded vertical position) has long represented a paradigmatic case for the theory of sexual selection by female choice. However, even after 150 years, the problem of the evolution of the peacock’s long tail remains unsolved—we still do not know the basis (target) of female choice in peafowl. Darwin (1859) recognized the peacock’s long tail as a problem for his theory of evolution by natural selection as he conjectured it was too long to be of adaptive use to the animal; therefore, it was maladaptive. Unlike in other animals, where a sexually selected trait may be directly under female choice selection, there are three aspects of the peacock’s tail: length, and structural complexity, and “beauty.” By beauty here we are concerned with the structural arrangement and the color of the eyespots, which adds to the overall beauty of the peacock’s tail. Darwin chose to focus on the beauty of the tail and supplied an explanation through sexual selection: that females may prefer to mate with males who possess more beautiful and elaborate tails (Darwin, 1871). It was thought that this reproductive advantage enjoyed by males with more elaborate tails would compensate for any loss of male fitness such as reduced survivorship due to predation. This explanation sets the stage for research to focus on elucidating how females assess “beauty” or attractive traits (in peacocks and other birds) in their choice of mates (Andersson, 1994).

Two key requisites for evolutionary theory, be it via natural or sexual selection, are variation and heritability. If the peacock’s train is a target of female choice, then there must be genetic or phenotypic variation in female preference that would directly or indirectly depend on variation in the peacock’s train morphology (size, shape, and/or coloration). Although female choice is widely assumed to be responsible for the evolution of the peacock’s tail, research on this matter has produced mixed results to date. While several researchers have shown a relationship between train features and mating success (Petrie et al., 1991; Petrie and Halliday, 1994; Yasmin and Yahya, 1996; Loyau et al., 2005a; Harikrishnan et al., 2010), a comprehensive 7-year re-evaluation by Takahashi et al. (2008) found no evidence of train morphology (either train length or eyespot number) influencing the number of copulations a male achieves. More importantly, this re-evaluation uncovered little variation in train morphology across populations, thereby calling into question the idea that the peacock’s train is a target of female choice (however, see Loyau et al., 2008). Similarly, in another important study based on close observation of eyespot distribution on the train, Dakin and Montgomerie (2011) found that males over the age of four generally produce between 165 and 170 eyespots before the onset of the mating season. They found little variation within and between populations in terms of eyespot number and noted that what little variation existed seemed to be due to extrinsic factors. Dakin and Montgomerie (2011) rather intuitively suggested this lack of variation in train morphology possibly reflects developmental constraints. There may be variation in eyespot number between growing males (Manning, 1989) but the number in adult males appears invariant (Manning, 1989; Takahashi et al., 2008; Dakin and Montgomerie, 2011).

While much work has been done to investigate the basis of female choice, less work has addressed the structure of the trait, i.e., the train itself. The peacock’s train is a complex structure, with the upper train coverts comprising a diverse variety of feather types, each varying in structure, iridescence level, color pattern and symmetry (see Lillie, 1942 and Manning and Hartley, 1991 for details). The complexity and the bilateral symmetry of the train (Figure 1) can be appreciated by connecting the eyespots in any direction (see Figure 2A). To achieve this remarkable symmetry in a fan formation, the feather follicles must develop in a specific arrangement at their origin, i.e., on the uropygium. Upon failing to find significant variation within and between populations in terms of eyespot number, Dakin and Montgomerie (2011) inspected the outer surface of a male peacock’s uropygium and found a constant number and uniform arrangement of feathers, which suggests that train feather development may be anatomically determined in all males. This suggests that the extraordinary symmetry of the train (Figs. 1, 2) is dictated by developmental plans which may not allow for intrinsic variation in train morphology.

**Figure 1.**
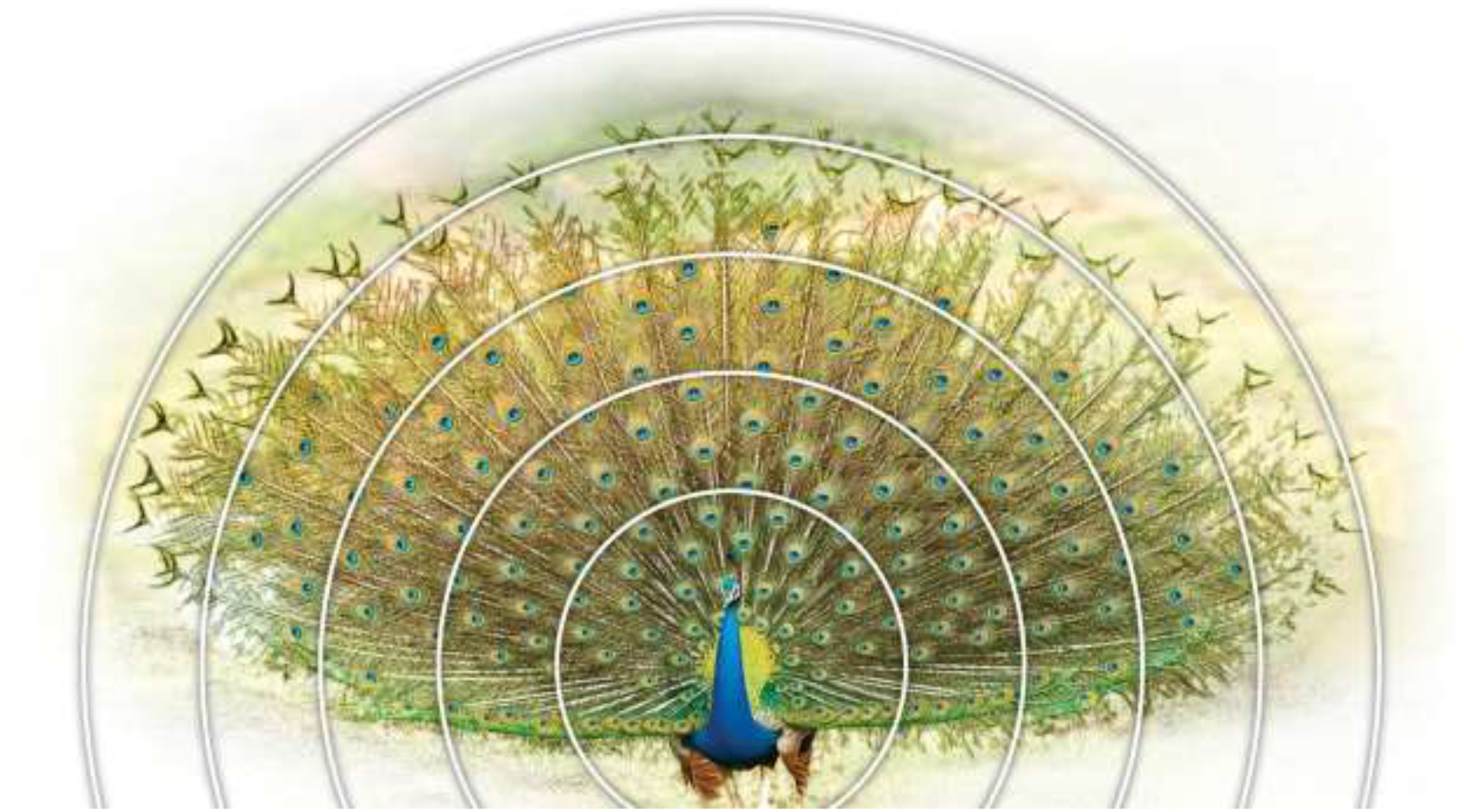
A picture of a peacock tail showing the symmetry of the train and eyespots. Concentric circles show spaced-out eyespots towards the periphery (picture from Wikipedia/ ThiminduGoonatil lake from Colombo, Sri Lanka).

**Figure 2.**
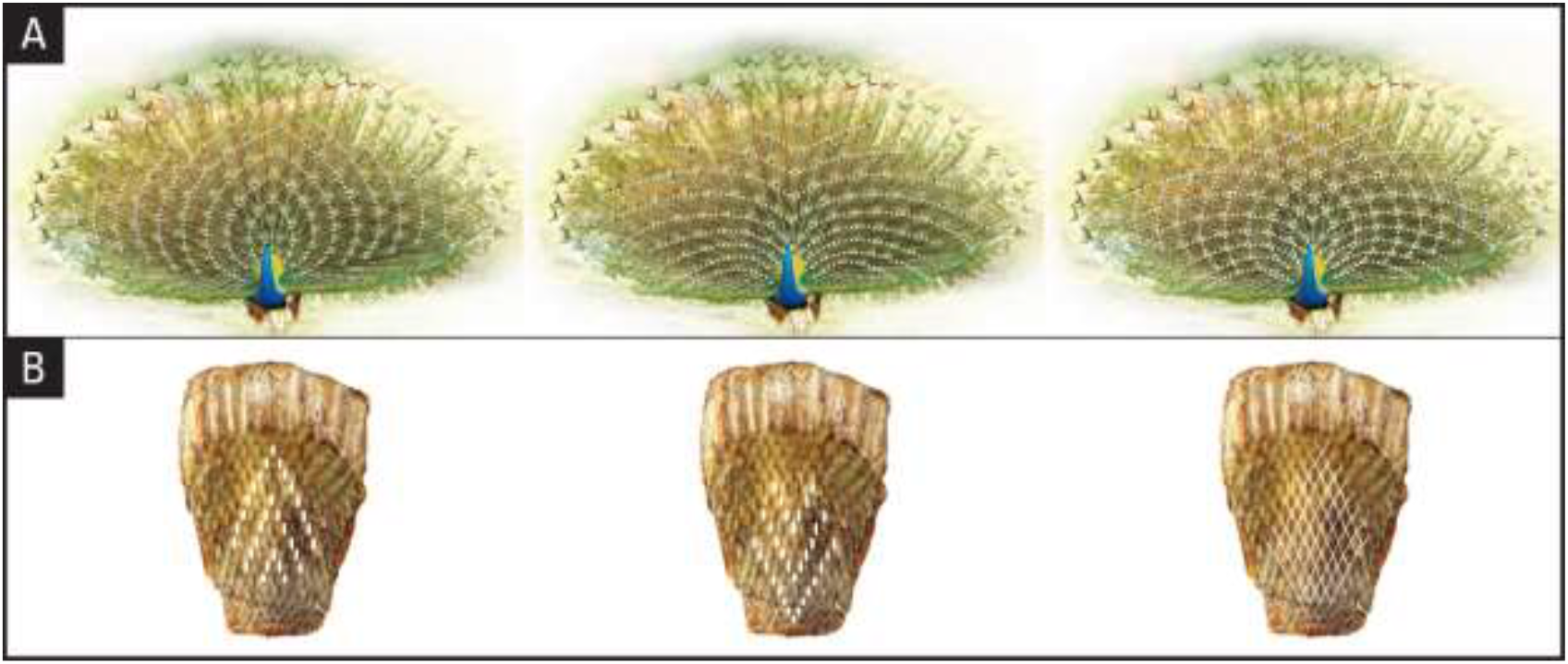
Geometrical designs and the symmetry of the train are indicated by dotted lines (**A**) and the position of follicles on the anchor plate (**B**). (Peacock picture from Wikipedia/ThiminduGoonatil lake from Colombo, Sri Lanka).

To explain the remarkable symmetry of the structural arrangement and iridescence of the train and to understand the nature of variation in eyespot number we investigated the anatomical development plan of peacock train morphology. We uncovered developmental and anatomical evidence demonstrating that the nature of bilateral symmetry in the development of the peacock’s upper-tail covert feathers or train feathers) would preclude genetic variation in the number of individual eyespots. Accordingly, the only source of intrinsic variation in feather and eyespot number would be between age classes, arising from the age-dependent addition of new rows of feathers. On the other hand, feather length—being a quantitative trait—is expected to show variation within as well as between age classes. Thus, all intrinsic between-individual variation in eyespots may be related to animal age, which may reach a developmentally pre-determined limit in all animals (Dakin and Montgomerie, 2011). This would explain why researchers have not found significant (genetic) variation in eyespot number in natural populations (Takahashi et al., 2008; Dakin and Montgomerie, 2011).

In this report based on observations on museum specimens of peacock tails, we show that a simple zigzag pattern of feather formation giving rise to hexagonal arrangement of eyespots uniquely determines the symmetry, complexity, and beauty of the peacock’s tail. Hexagonal arrangement of identical circles is the most efficient form of packing, and it was shown to apply to the origin of feathers in birds (Sengel, 1976). By beauty, in this manuscript, we only mean the complexity of the eyespot distribution and the color patterns and do not want to confuse with everyday and wider meaning of the word beauty. Second, we show that eyespot feathers originate in alternate, zigzagging rows of 10/11 annually, making the total number of eyespots an intrinsically determined trait. Third, we argue that both feather number (Manning, 1989) and feather growth are asymptotic functions of age that—when considered with the male vigor which must peak at reproduction time and then decline—may explain the contradictory results reported between different studies (Loyau et al., 2008; Takahashi et al., 2008). We discuss the implications of these results, especially the lack of genetic variation in eyespot number, for sexual selection theories and how alternative explanations are required to explain the basis of female choice and the evolution of the peacock’s elaborate train. We propose a multimodal model of female choice based on male’s train size, vigor, and beauty that aligns with recent calls for more inclusive and multimodal perspectives to understand how sexual selection operates (Mitoyen et al., 2019), as it highlights the multiple factors which may jointly determine the female choice of best mates. We propose that many conflicting results can be explained by assuming that females do not base their mate choice on eyespot number or train size alone but on a complex trait combining both that may act like a bright “moving mural.”

## 2. Materials and Methods

### 2.1. Peacock specimens

The data presented in this report are based on observations of the tail structures of museum specimens kept at the Royal Ontario Museum, Toronto, Canada, and the American Museum of Natural History, New York, USA. Only specimens in good condition were included in this study. A total of 21 samples of blue fowl (*P. cristatus*), including two albinos and two hybrids (between strains originating from Bangkok and Cameroon), were examined. The wild blue fowl samples originated from India (N = 3), Sri Lanka (N = 2), Kenya (N = 1), or were captive samples originating from the United States (N = 4 blue, N = 2 albino). The remaining 11 samples were of unknown origin and included two hybrids. We also had access to seven green fowl (*P. muticus*) samples at the Natural History Museum, which originated from Malaysia (N = 4), Bangkok (N = 1), or were of unknown origin (N = 2). Wherever dates were noted, most of the field samples were collected during the early 1900s. Conversely, the captive samples were collected as recently as 1942.

### 2.2. Data collection

We counted the number of eyespots, eyespot feathers, and fishtail feathers displayed by the included specimens (Figure 3). In the cases where an eyespot was missing due to damage, we counted it as if it were present to estimate the total number of eyespots. Very few of the samples had a uropygium that was in good condition. Of the museum specimens six had intact uropygia in *P. cristatus* and one in *P. muticus*. Wherever we were certain that the integrity of the sample had not been compromised, we counted the rows of feather imprints on the uropygium (Figures 2B, 4). For raw data, see Singh and Jagadeeshan (2021).

**Figure 3.**
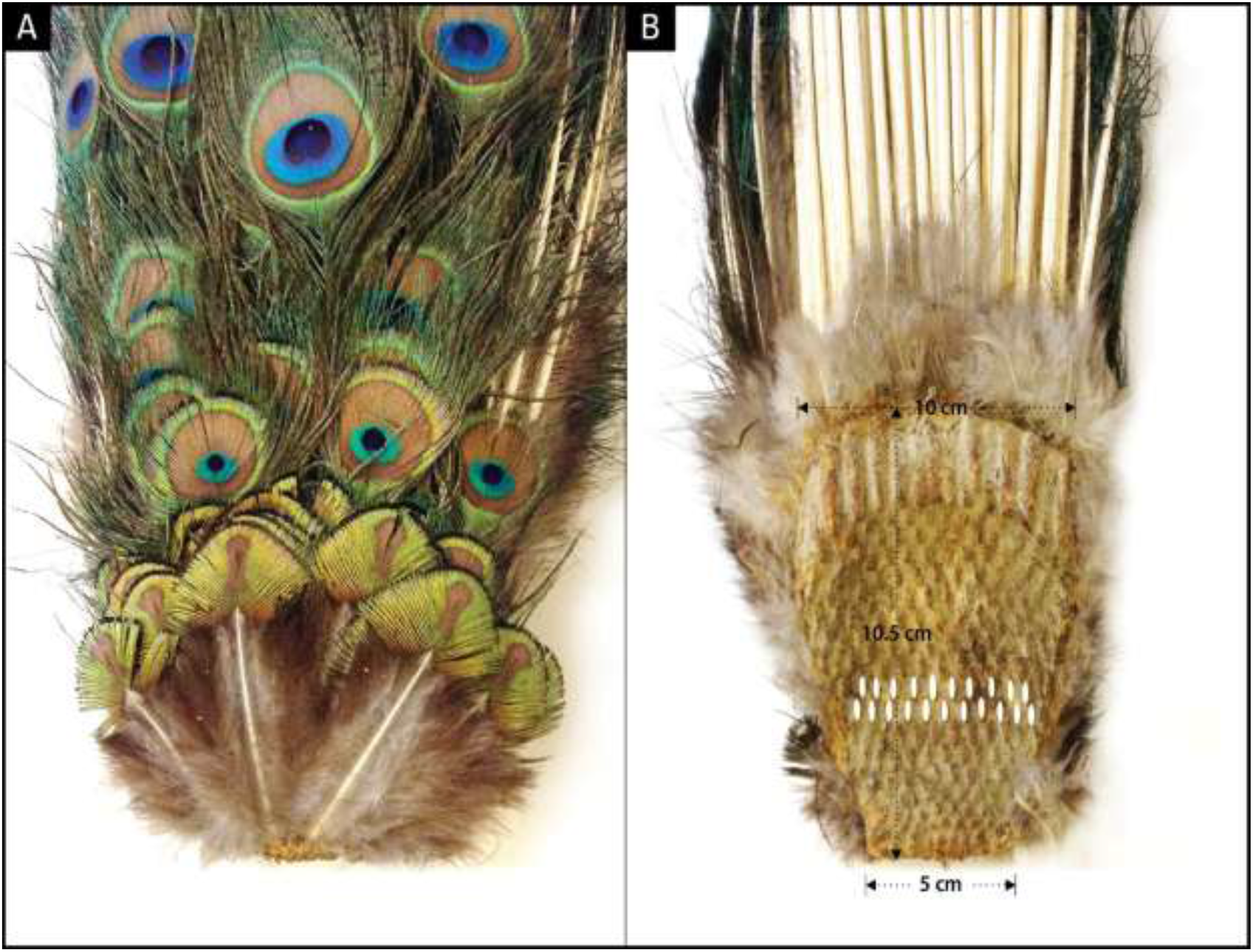
A: Dorsal side of the anterior portion of the peacock train showing the newest rows of feathers at the bottom; **B**: Ventral side of the Train showing the uropygium, depicting alternating rows of 10 or 11 feather cell imprints. The smallest feather row at the bottom of panel **A** corresponds to the smallest row of follicles at the bottom of panel **B**. The top 3-5 rows depending on the age of the animal give rise to fishtail feathers and the rest eyespot feathers.

### 2.3. Simulating train expansion

To illustrate the eyespot symmetry exhibited by trains in displays (Figure 5), we made use of Adobe Illustrator’s built-in features (Adobe Photoshop and Adobe InDesign) provided by Graphic Designers’ services at Media Production Services, McMaster University. Briefly, Adobe Illustrator was used to map concentric circles following eyespot distribution on the train in the display by using Figure 1 as a template. These sets of concentric circles were used as the basis for text paths to determine how eyespots may distribute themselves upon the unfolding of the train for display.

**Figure 4.**
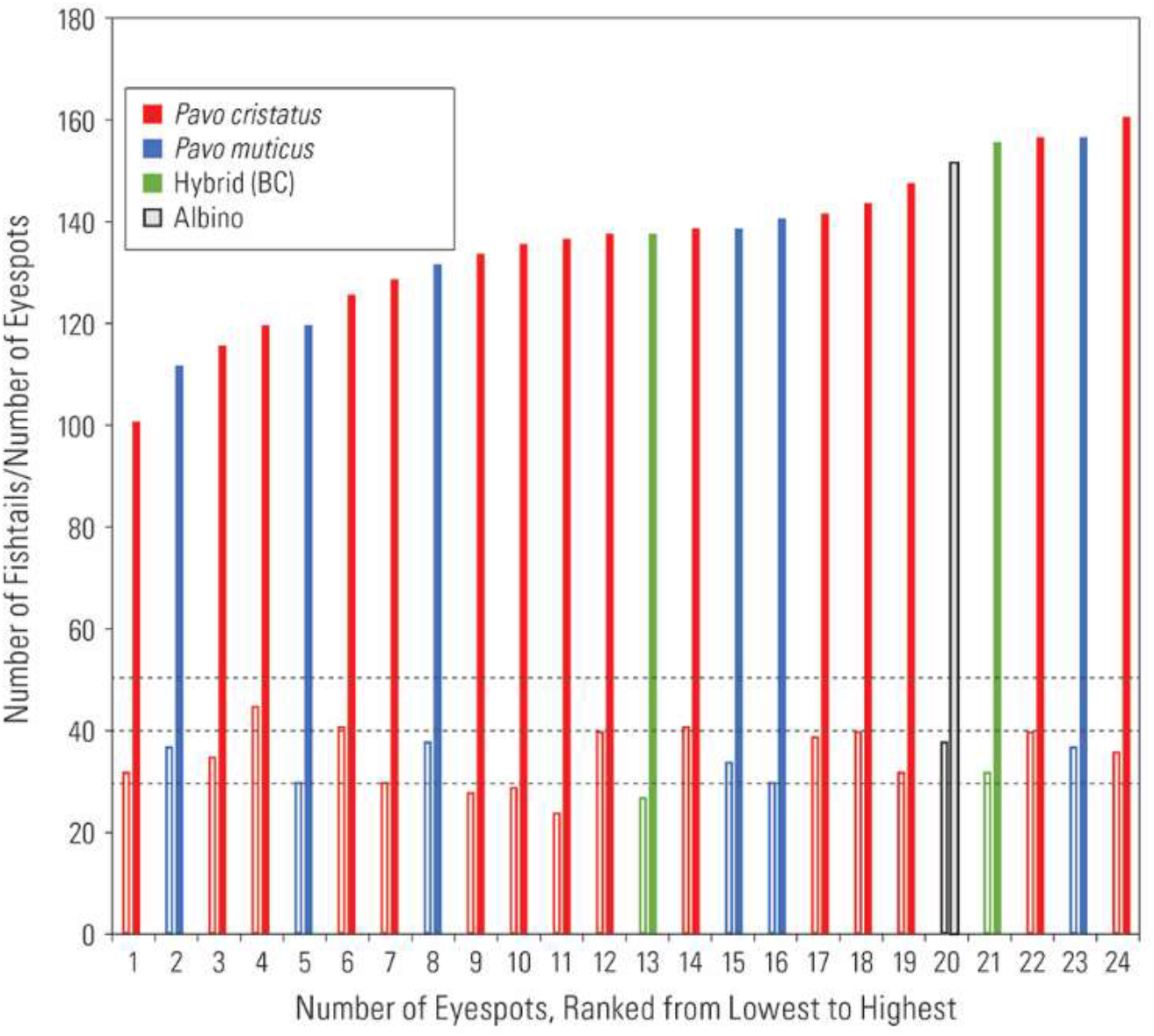
A graph showing independence of eyespot and fishtail feather numbers. Dotted lines represent number of fishtails (30+, 40+, 50+) consisting of 3, 4, and 5 rows of feathers (details provided in the text). Closed bar: eyespot feathers; open bar: fishtail feathers. The hybrid (BC) is between Bangkok and Cameroon.

**Figure 5.**
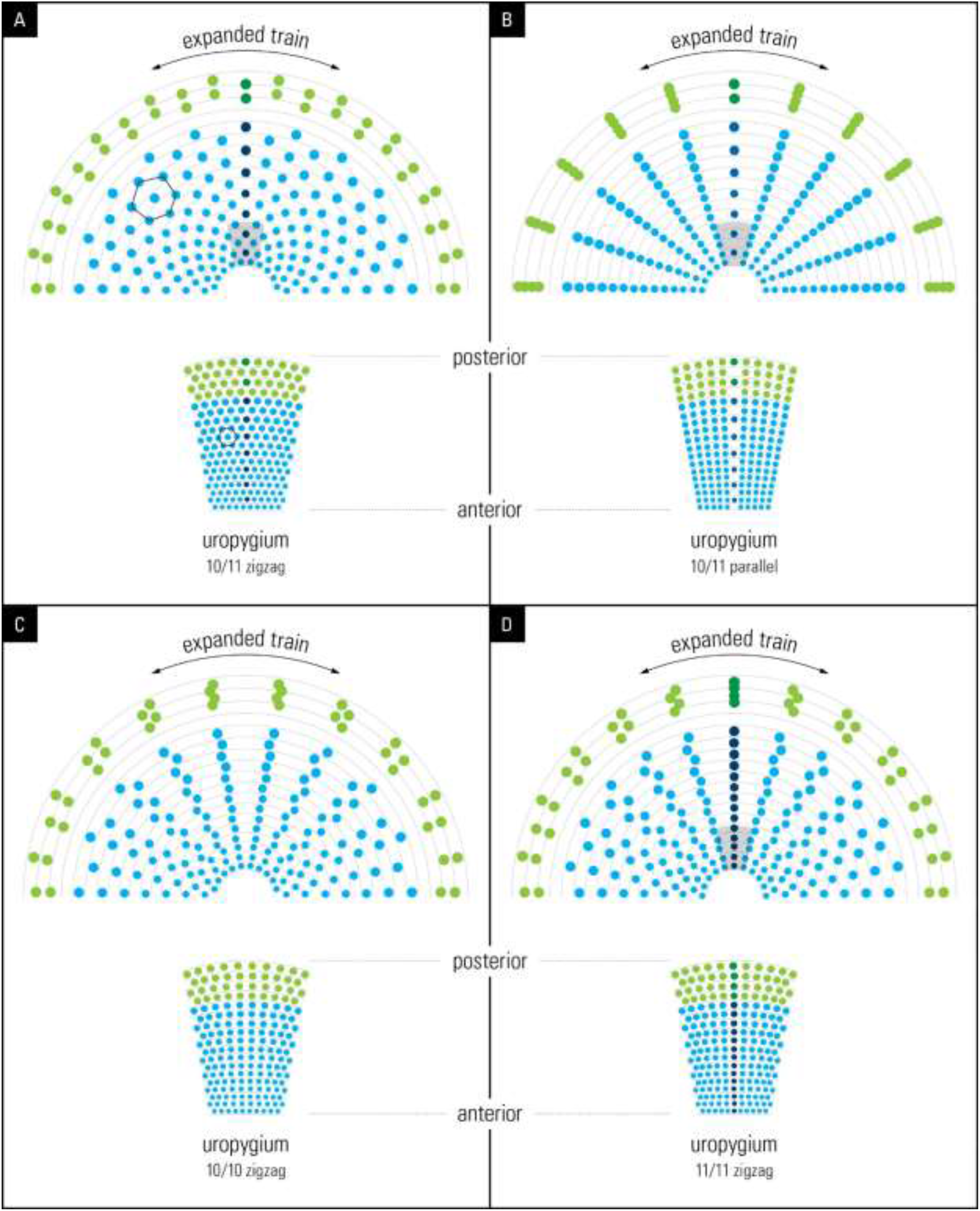
Illustrations from graphic simulations of eyespot symmetry showing the layout of feathers on the anchor plate (below) and the resulting eyespot patterns produced by a bilateral symmetrical train (above). The outer spots (green) represent fishtail feathers. **(A)** 10/11 zigzag pattern; **(B)** 10/11 parallel pattern; **(C)**: 10/10 zigzag pattern, **(D)** 11/11 zigzag pattern. Compare the hexagon cellular packing on the uropygium and the expanded train in **A** with that seen on the animal’s train in Figure 1(for details see the text).

To simulate and illustrate the train’s eyespot distribution, we used the following observations and/or assumptions: (1) Based on our overall observations (7 *P. cristatus*, and 1 *P. muticus*), we concluded that each cell on the uropygium (Figure 1B) represents the base of a corresponding feather (attempting to determine direct one-to-one correspondence would have required damaging the specimens); (2) we used a row of 10 or 11 dots in the shape of the (oval) uropygium to represent train feathers but we used a flat and not a convex surface as the latter was not possible. In the result section we discuss why using a convex surface in the simulation would not have changed the outcome; (3) In line with our observation (Figure 1), we used an arbitrary constant distance between spots and we progressively increased the size of the dot spots between rows, from the bottom (newest) to the top (oldest) feathers, to represent the size of the eyespots; (4) We assumed that each dot represents an eyespot at the end of the feather which is supported by our observation; and (5) we only simulated 2-dimensional position of eyespot distribution and not their three dimensional distribution on the train surface. We simulated four spatial feather arrangements: 10/11 zigzag, 10/11 parallel, 10/10 zigzag, and 11/11 zigzag (Figure 5). The feather grid was expanded bilaterally symmetrically to mimic the animal’s train expansion. It is important to point out that the simulated surface shown in Fig 5 represents the front face of the train.

## 3. Results

### 3.1. The origin of bilateral symmetry in the peacock’s train

To uncover the anatomical development plan that determines the bilateral symmetry of eyespots on the train, we examined the uropygium (N = 6) of museum specimens. The uropygium is a fleshy and bony structure at the posterior extremity of a bird’s body that supports the tail feathers. Tail feathers are attached to the dorsal (convex) side of the oval-shaped uropygium, which serves as an anchor plate (Figure 3). The ventral (concave) side of the anchor plate exhibits several interesting features: 1) It has basal impressions of tightly-packed, parallel rows of feather follicle insertion points; 2) The feather follicles grow progressively larger from the anterior (bottom, younger) to the posterior (top, older) end and are laid out in sequential but alternating rows of 10/11, corroborating the numerical feather arrangement of 10/11 first observed by Dakin and Montgomerie (2011), except that while we had expected them to be in parallel rows based on our interpretation of Dakin and Montgomerie’s work (Fig 1A in Dakin and Montgomerie, 2011), but they were in a “zigzag arrangement” (see Figure 3, right); 3) The posterior feather follicles radiate in an arch, mimicking the fan formation shape of the train in the display. The convexity of the anchor plate makes the train a three-dimensional structure (i.e., an oval trapezoid-shaped dish), such that the feathers are projected outwards at different angles and lengths. Notably, the anchor plate itself appears to be a product of continuous growth and the sequential addition of new rows of feathers each year. The one-to-one correspondence between the pattern of the feather follicles (size, alternate arrangement, and progression from anterior to posterior) on the ventral side of the anchor plate and the symmetrical positions and size of the feathers on the dorsal side is unmistakable and is supported by our observations (Figure 2).

### 3.2. Variation in eyespot number

To determine whether any variation exists in the anatomical development plan found on the uropygium that would affect the number of eyespots, we examined several specimens from the Royal Ontario Museum and the American Museum of Natural History. The results of this analysis, based on a limited number of specimens in good shape, are summarized in Table 1 and Figure 4. First, the number of feather rows, as determined by the number of follicle rows on the uropygium, varied from 17 to 19 (N = 7), whilst the anterior two to three rows had minor eyespots. Notably, this result is consistent with the findings of Dakin and Montgomerie (2011). The anterior-most rows of eyespots appear minor due to their slow growth and maturity. If one or a few rows of feathers are added each year, then based on the peacock’s longevity of about 20 years in nature (Lindstedt and Calder, 1976) the number of eyespots can reach over 200; however, the late-aged, developmentally immature eyespots (small size and lack of full coloration) will remain insignificant in their effect on female choice.

**Table 1.**
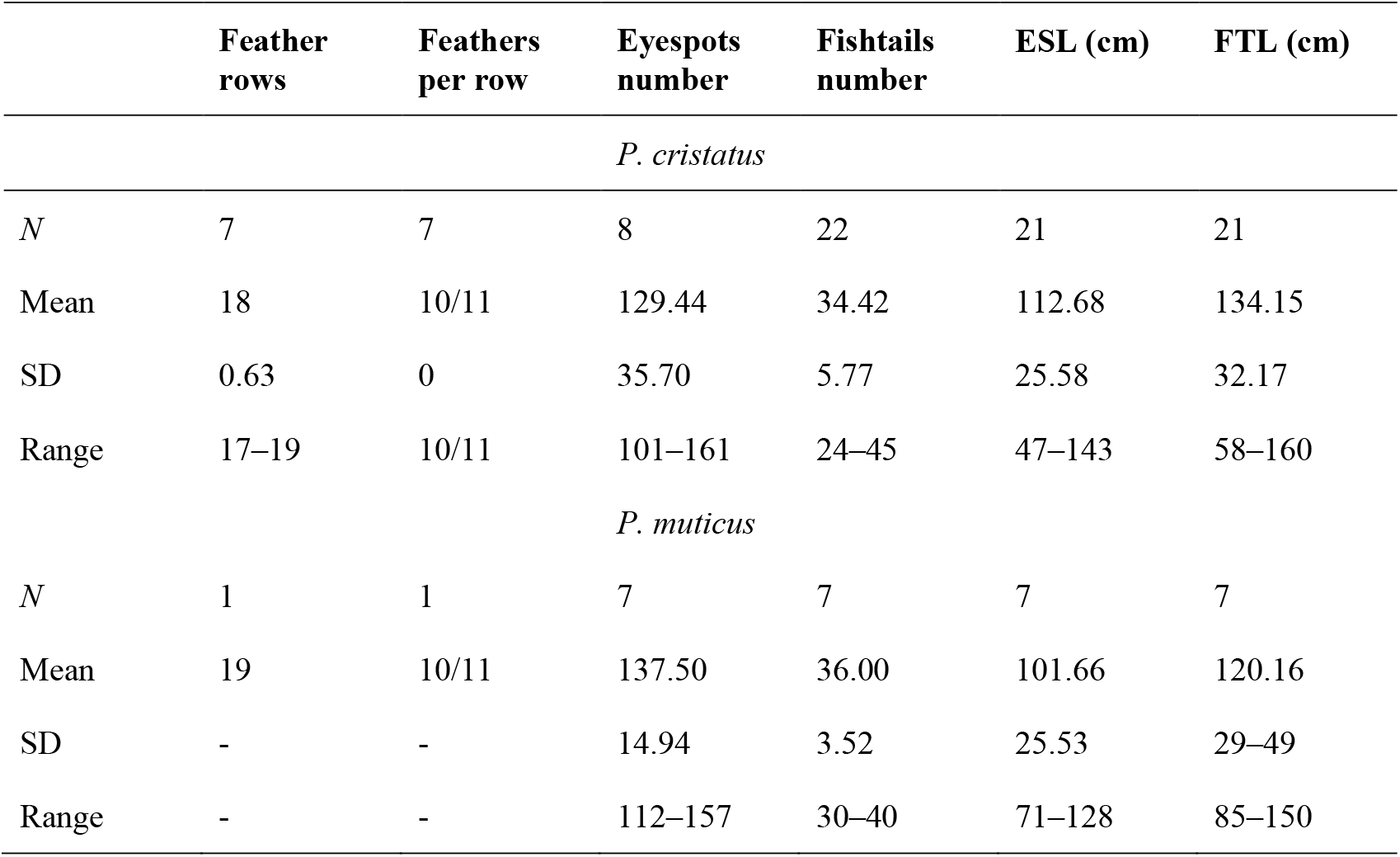
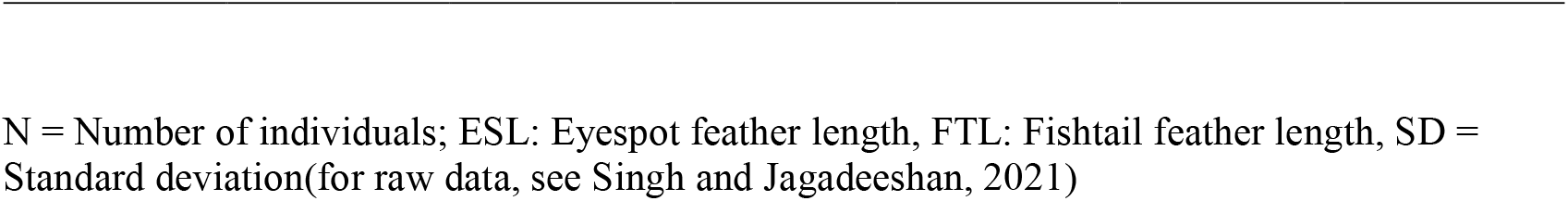
Variation in peacock train morphology between *P. cristatus* and *P. muticus*.

|  | Feather rows | Feathers per row | Eyespots number | Fishtails number | ESL (cm) | FTL (cm) |
| --- | --- | --- | --- | --- | --- | --- |
| *P. cristatus* |  |  |  |  |  |  |
| *N* | 7 | 7 | 8 | 22 | 21 | 21 |
| Mean | 18 | 10/11 | 129.44 | 34.42 | 112.68 | 134.15 |
| SD | 0.63 | 0 | 35.70 | 5.77 | 25.58 | 32.17 |
| Range | 17–19 | 10/11 | 101–161 | 24–45 | 47–143 | 58–160 |
| *P. muticus* |  |  |  |  |  |  |
| *N* | 1 | 1 | 7 | 7 | 7 | 7 |
| Mean | 19 | 10/11 | 137.50 | 36.00 | 101.66 | 120.16 |
| SD | - | - | 14.94 | 3.52 | 25.53 | 29–49 |
| Range | - | - | 112–157 | 30–40 | 71–128 | 85–150 |

N = Number of individuals; ESL: Eyespot feather length, FTL: Fishtail feather length, SD = Standard deviation(for raw data, see Singh and Jagadeeshan, 2021)

Second, the zigzag arrangement of feathers appeared as an invariant trait in our sample (Table 1). While this one-to-one correspondence between clearly visible follicle insertion patterns and train morphology may be expected, our data suggest that the developmental plan does not permit random (one feather at a time) variation in eyespot number (Table 1).

Third, the number of fishtail feathers varied around the modal number of 30, 40, or 50 (N = 21), which is what we would expect if fishtail feathers were produced by three, four, or five rows of feather follicles during development, respectively (Figure 4). Depending on whether the first row is made of 10 or 11 feathers, the number of fishtails would vary from 31–32, 42, or 52–53. In our museum samples, the highest number of fishtail feathers observed was around 40. However, Figure 1 indicates that the number of fishtail feathers can exceed 50.

Fourth, the number of eyespot feathers varied from 101 to 161 (N = 21), not counting the minor feathers. This variation is attributable to individual variation in terms of age, time of year at capture, and damage from prior handling at the museum. Fifth, number of eyespot feathers and feather length are independent traits that vary asymptotically as a function of age (data not shown). This is consistent with Manning’s (1989) finding that eyespot number increases during the first 4 to 7 years of the animal’s life and thereafter increases more slowly or remains effectively constant.

### 3.3. Interspecies variation: *P. cristatus* vs. *P. muticus*

We also had access to a small number of *P. muticus* (green peacock) specimens (*N* = 7) from Southeast Asia (Table 1; Figure 4) for comparative analyses, which could provide clues as to whether peacock train morphology reflects a similar developmental pattern across taxa or whether sexual selection and other environmental factors have resulted in different trajectories of train morphology. In the green peacock, the number of fishtails ranged from 38 to 39 (around the expected mode of 42), while the number of eyespots ranged from 115 to 154.

Data from both species appeared to be homogenous (Figure 4). One green peacock specimen was in pristine condition, which enabled us to assess the anatomical development pattern of feather follicle rows in the uropygium. As per the blue peacock, feather follicle rows in the green fowl followed the same alternating 10/11 arrangement, suggesting that this developmental arrangement may be an invariant feature across peafowl species. Interestingly, similar alternating patterns of train feather arrangements are also found in the wild turkey, *Meleagris ocellata*, a member of the same family (Phasianidae). This observation entertains further research on whether the alternating pattern of feather arrangement might be common across other members of the family and thus an invariant developmental trait. N = Number of individuals; ESL: Eyespot feather length, FTL: Fishtail feather length, SD = Standard deviation(for raw data, see Singh and Jagadeeshan, 2021)

### 3.4. Speciation *in-silico*: The origin of the train’s symmetry and complexity

We investigated the significance of the arrangement of feathers in rows of 10/11 and their “zigzag” alignment (see Figure 2). To achieve this, we used graphic design software to reconstruct the unfolding of the peacock’s open train based on the developmental layout of the feather follicle insertion points observed in the anchor plate (Figure 5, left). For simplicity, we assumed that the feather follicle insertion points were eyespots without the feather stalks. Using Adobe Illustrator, we modeled the fan formation in “display.” We subjected the eyespot distribution on the anchor plate to a uniform, bilaterally symmetrical force to spread the eyespots to the left, right, and upward, thereby pushing them radially and uniformly outward in a curved semi-circular space to mimic the fanning pattern of the train during display. Remarkably, our reconstruction yielded an eyespot distribution that is very similar to that observed on the peacock train (compare Figure 5A to Figure 2), with five feathers on either side of the mid-feather in each row. As expected, we found a one-to-one correspondence between the condensed, developmental–anatomical layout of the feather follicle insertion points on the uropygium and the symmetrical distribution of eyespots on the train. It is important to point out that the zigzag arrangement of cells, when spread out uniformly, gives rise to the well-known hexagon cellular packing (compare Figure 5A and Figure 1). We did not simulate a convex surface as this was not possible, however doing so would not have changed the distribution pattern (hexagonal) of eyespot as we were simulating 2-D image of eyespots distribution as seen from front and not their 3-D distribution patterns. In other words, the simulation results presented in Figure 5 show what the imprint/picture would look like if an expanded train were compressed on a flat surface. This is what we would see from a distance.

We further explored what the eyespots distribution pattern would be like if the feathers were arranged in 10/11 parallel rows instead of the zigzag arrangement that we observed in this study. Notably, the train eyespot pattern that we obtained is remarkably different from the 10/11 zigzag pattern. Our reconstruction of a parallel arrangement yielded a palm-leaf-like pattern that fanned out in parallel rows of eyespots rather than the pattern observed in a peacock’s train on display (Figure 5B). While both patterns are striking, the 10/11 zigzag arrangement yields a denser and uniformly symmetrical arrangement of eyespots, as seen in the animal.

Because the above results raised the question of why 10 or 11 rows were observed, we simulated a 10/10 parallel and 11/11 parallel pattern; the results are shown in Figures 5C and D. Although the 10/10 and 11/11 patterns were like the 10/11 parallel pattern, they showed certain interesting differences. While the feathers lined up in equally spaced, straight lines over the entire span in the 10/11 parallel pattern, the 10/10 and 11/11 parallel patterns resulted in the equal spacing of individual eyespots towards the end of the train, with wavy lines in the center. It is obvious that it was not the zigzag pattern itself but the zigzag pattern of 10/11 that gave rise to the uniform distribution of eyespots on the train. The 10/11 parallel pattern may not be developmentally possible.

### 3.5. A rudimentary model of eyespot development

Feather is an important non-embryonic model of animal development (for a recent review see Chen et al., 2015), and much research has been done both anatomical and molecular involving the role of genetic and epigenetic mechanisms (Jiang et al., 1999, 2004; Maini et al., 2006; Chen et al., 2015). Feathers develop from feather follicle buds on the surface of the epidermis (Prum, 1999; Chen et al., 2015). Pigmentation based colors in plants and animals can be explained by local variation in gene expression. In birds, movement of melanocytes up the hollow core of the feather is said to control the variation in plumage color (Prum and Williamson, 2002; Chen et al. 2015). In “structural color,” such as in peacock feathers, the colors are the result of light reflection from the modification and anatomical diversity of the barbules’ photonic structures (Zi et al., 2003, Freyer at al. 2018).

We were motivated by the superficial but interesting structural similarity between the iridescence patterning of the eyespots and that of the body plan of the whole animal with a fully expanded train. The entire peacock resembles one giant eyespot in the sense that it boasts a deep-blue body surrounded by a dense zone of bluish-green eyespots, which corresponds to an individual eyespot with its deep-blue inner circle surrounded by radiating oval rings of mixed colors. The same pattern of blue can be observed in the peacock’s crest crown also (data not shown). The peacock’s crest consists of a row of silky threads that are a few centimeters in height and end in a feathery crown; they have been shown to be involved in communication (Kane et al., 2018).

We hypothesized that the rings of eyespots may be determined by the same cellular plans as the color patterning of the whole animal. Such deep developmental plans affecting structural color in different traits may be expected but how female choice may have evolved in relation to the eye color is not known. We took advantage of the similarity between the rings on the individual eyespots and the spatial eyespot rings traced on the train and modeled backward from the eyespot to feather follicles and back to the eyespot (Figure 6). Starting from each eyespot (Figure 7a) we inferred colored rings on the train following the pattern seen in one eyespot (Figure 6b) and from there on the follicle rows on the uropygium (Figure 6c). We inferred that the five feather follicles on each side of the central feather make an inverted palindrome and correspond to the five different structural colors seen in the eyespot (Figure 6d). From this we inferred follicles growth and epidermal invagination giving rise to a concentric ring of structural material (stem cells) inside the feathers which expand and create the ring pattern on the eyespot. Cellular invagination is a well-known process of embryology (Rauzi et al., 2013; Pearl et al., 2016) but we stress that we are basically connecting dots from eyespot distribution on the train to the concentric rings of individual eyespots and it must remain speculation until further investigated. This structural developmental plan underscores the remarkable similarity between the color patterns of the whole animal, the eyespot, and even the head crest. If experimentally proved, the deep-seated developmental plan of feather development may explain the invariant nature of the eyespot number across taxa.

**Figure 6.**
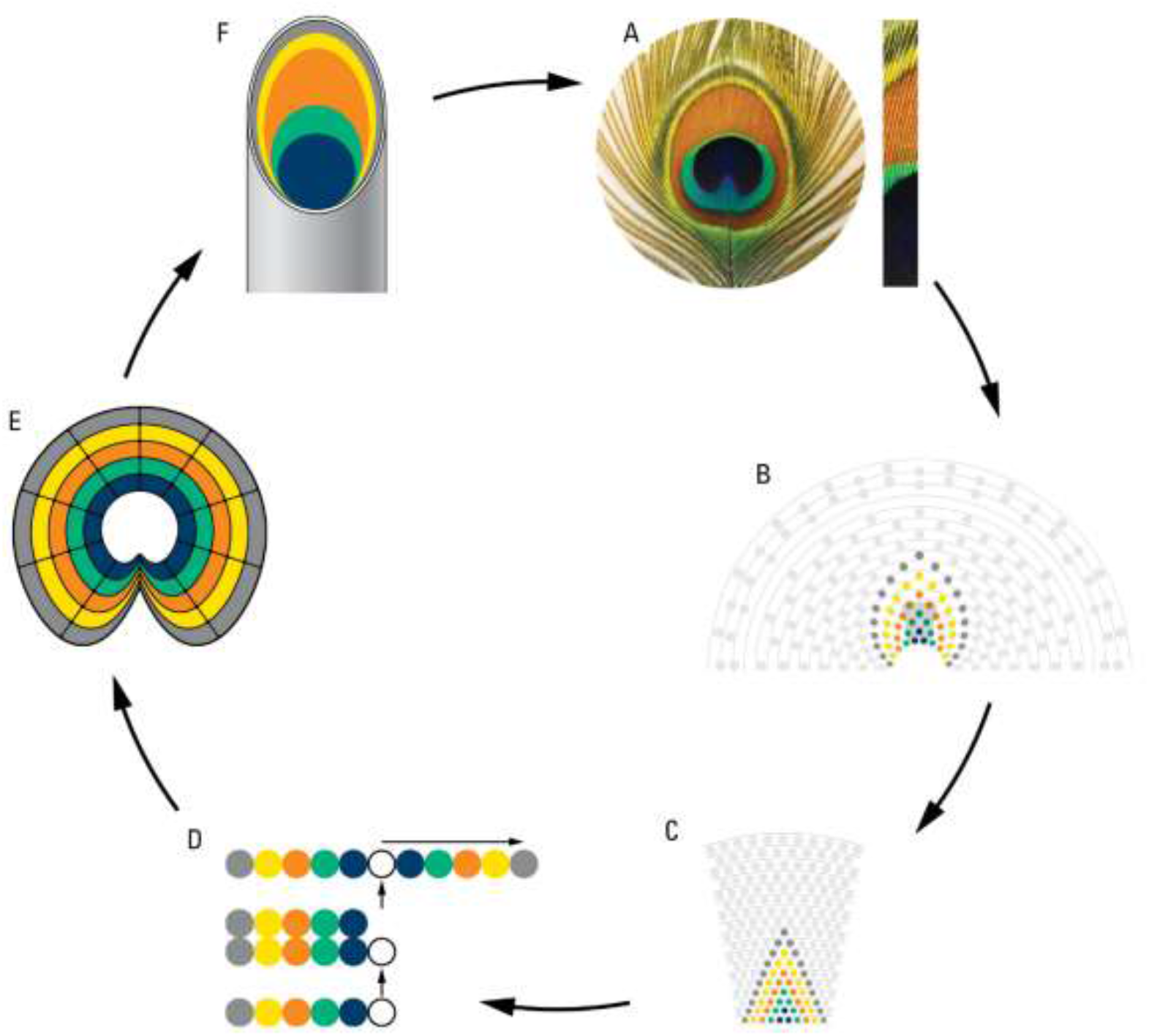
A diagrammatic representation of a theoretical plan of eyespot development as deduced from eyespot to follicles, and back to eyespot. **A**. an eyespot and a bunch of eyespot fine feathers showing their multi-colored structures; **B**. modelling of eyespot rings using rings observed on the train; **C**. projection of train rings on the uropygium; **D**. cellular processes showing origination of 5 structurally different feather follicles, and their inverted tandem duplication; **E**. cell growth and invagination of a concentric ring of structural material giving rise to eyespot; **F**. a peacock feather showing internal component of eyespot fine feathers (for details see the text).

## 4. Discussion

Mating success of peacocks has been shown to be affected by eyespots and train size, which appear to be correlated, but it has not been possible to determine if one or both affect mating success directly (Petrie et al., 1991; Petrie and Halliday, 1994; Yasmin and Yahya, 1996; Loyau et al., 2005a; Harikrishnan et al., 2020). Reducing the number of eyespots has a negative effect on mating success (Petrie and Halliday, 1994); however, Dakin and Montgomerie (2011) showed that females were insensitive to removal of as many as 20% of the eyespots before responding to the reduction. Petrie and Halliday (1991) surmised that “eyespot number or *something closely related to it* is important for female choice (emphasis added).” Could that “something” be the train length? In the following, after showing that eyespot number and train length are intricately related and that their roles as cues for female choice are largely inseparable—and, in addition, that male vigor is an important factor mediating the relationship between males and females—we present a multimodal (Anderson, 1994; Candolin, 2003; Bro-Jorgensen, 2010; Mitoyen et al., 2019), two-stage model of female choice based on a complex trait combining both train length and eyespots that may act as a mate attractant from a distance and as a mark of mate quality, vigor and beauty from close proximity.

### 4.1. The zigzag feather pattern explains the train’s symmetry and complexity

The zigzag pattern of feather formation observed from the peacock’s uropygium was able to explain both the hexagonal eyespot symmetry on the Train and as well as to the unique eyespot rings pattern within eyespots.

The primordial feather follicles arising on the uropygium in rows of 10/11 are arranged in a zigzag manner. It is this zigzag arrangement of 10/11 that creates the complex hexagonal arrangement of eyespots in the peacock’s tail, and thus its complexity and beauty (Figure 5, upper left). Hexagonal arrangement is a well-known feature of cellular arrangement in developmental biology (Sengel, 1976; Classen et al., 2005). The same rule of zigzag arrangement between feather follicles that leads to the complex distribution of eyespots on the train also applies to cellular arrangement of cell lineages within feathers, giving rise to rings on the eyespot, which, in combination with the different structural -iridescence colors, imparts the eyespot its beauty. The qualitative annual growth of eyespot number in rows of 10/11 and the quantitative growth of train length and, essentially, of the former’s dependence on the latter for its presentation and display to females may make the two traits inseparable in their effects on female choice.

### 4.2. Pattern of feather development constrains eyespot variation pattern

#### 4.2.1. Bilateral symmetry bars random variation in eyespot number

The anatomical arrangement and bilateral symmetry of eyespot feathers preclude random genetic variation by addition of single feather at a time and the only apparent source of variation in eyespot number is the annual addition of new feather rows as the animal grows. Thus, as Takahashi et al. (2008) and Dakin and Montgomerie (2011) found, most males produce approximately 165–170 eyespots, and the small amount of eyespot number variation that exists within and between populations is due to extrinsic factors such as breakage or damage from predation. The actual number of eyespots can exceed 200, as suggested by the number of feather rows on the uropygium shown in Figure 3, but many of them would be too small to be visible through photography or to be effective to illicit female response. The idea that the maximum eyespot number in adult animals is invariant within and across populations is consistent with the annual addition of new rows of feathers, which occurs rapidly at a younger age (∼4–7 years), while slowed growth or complete cessation of new feather development occurs after a certain age (Manning, 1989). It is important to point out that the lack of variation in eyespot number applies to the lifetime total number and not to the population, which may contain eyespot variation arising from different age classes.

#### 4.2.2. Diminishing returns in fitness and asymptotic tail growth

After a male has reached his peak reproductive age and achieved the threshold train size to begin mating, his mating success will increase as a function of both the size of his train and his vigor in mounting a display. But with declining vigor with age and competition from younger males, the individual fitness function will plateau and decline despite large train size. At this point, the law of diminishing returns will commence, and each additional row of feathers will grow slowly, making them look minor, and exert a minimal effect on the feather length and fitness function. Moreover, limitations with respect to maximum eyespot number and the lack of correlation between train length and mating success may—as suggested by Takahashi et al. (2008)—indicate that these traits have reached a threshold. And since males do vary in mating success (Petrie et al., 1991; Yasmin and Yahya, 1996; Loyau et al., 2008), this variation cannot be due to variation in eyespot numbers and other factors may be involved.

### 4.3. Implication for sexual selection theories

Female choice theory, in the case of the peacock’s tail, relies on the assumption that eyespot number and train length are correlated, and that female choice based on the former leads to a correlated increase in the latter. However, the results of this study, and others (Takahashi et al., 2008; Dakin and Montgomerie, 2011), suggest that eyespot number is an internally-determined, invariable trait, and thus cannot be the sole factor driving the evolution of the peacock’s long tail. Given that train feathers are rapidly added in rows of 10 or 11 during the first 4–7 years of a peacock’s life (Manning, 1989) to eventually reach 165–170 at adulthood, any variation in feather growth rate could be the basis of female choice if the size of the train were directly or indirectly a criterion of female mate attraction—or attention. As such, rapidly growing males may develop more elaborate trains with bigger size and more eyespots and thus become preferential targets of female choice during the peak period in the first 4–7 years of life, after which feather number may increase at a slower rate (Manning, 1989).

#### 4.3.1. A multimodal model of female choice and a hypothesis

Our study brings out three important results that have implications for how females decide on mates. First, the invariant nature of the total number of eyespots in adult males does not allow for variation in fully adult males for selection; the only available variation is that between age classes in a population. Second, eyespots and train length are not just correlated but are developmentally interconnected; it is the tail growth that pulls the eyespots up, spreads them out, and makes them visible to females. While eyespot brightness has been shown to affect male mating success (Loyau et al., 2007; Dakin and Montgomerie, 2013), we hypothesize that in deciding on mates, females may be using the size of the “eyespot-studded train” rather than the number of eyespots. This may explain why it has been difficult to separate the effect of eyespots from the effect of train size (Petrie et al., 1991). Finally, an important and hitherto neglected trait that is likely to mediate male–female interaction is male vigor. While eyespot number and train size are expected to grow asymptotically, male vigor must peak during mid-age and then decline. Variation in vigor means that a younger youthful male with fewer eyespots may outperform an older and bigger male with more eyespots (Petrie, 1993). The new trait, eyespot–train size, may either influence female choice directly through sensory capture (Kirkpatrick and Ryan, 1991) or indirectly through male vigor and beauty, but not solely based on eyespots (Mitoyen et al., 2019).

We can consider a two-stage model of female choice, in which females are attracted by male size or train height and sounds from a distance, and exercise choice based on eyespot beauty, vigor, and behavior from close proximity.

If females were making their choice based on the eyespot-studded train size and not based on eyespot number, an interactive model based on eyespot, train size and male vigor would be able to explain some of the contradictory results between different studies. It may explain, for example, why females were insensitive to removal of as much as 20% of the eyespots before showing any effect in their behavior (Dakin and Montgomerie, 2011).

### 4.4. Limitation and future work

The small sample size of museum specimens used in this study may raise concerns about the validity of our results. The most important results of this study, shown in Figure 5 and Figure 7, are qualitative and/or theoretical modelling, and are unaffected by sample size. Simulation is a form of hypothesis testing, but it lacks direct anatomical testing which is outside the scope of our (*Drosophila*) lab. The zigzag arrangement of feathers and its effect on the train’s symmetry and complexity (Figure 5) were simulated and shown to conform to observations from this work and at least the number and the symmetry of feathers is supported by previous work (Dakin and Montgomerie, 2011). The rudimentary model of eyespot development (Figure 7) would also need to be investigated by developmental work, which is also outside the scope of our lab. Future tests can involve (1) anatomical dissection of live animals to test the number and the arrangement of feathers as well as the correspondence between feathers and feather buds, (2) developmental work on young peacocks to test the addition of feather rows as well as feather growth as a function of age, (3) further work on the line of Dakin and Montgomerie (2011) to test the effect of varying eyespot numbers on female choice, and (4) work to test the effect of variation in male vigor as a function of age on female choice.

## 5. Conclusions

In this study we made three seminal observations that provide a new perspective on the evolution of the peacock’s tail. First, we showed that a zigzag pattern of feather formation affects the bilateral symmetry, eyespot complexity, and beauty of the peacock’s tail. Second, the same zigzag pattern, remarkably, can also explain the colorful rings of the eyespot. Finally, the zigzag pattern would preclude intrinsic variation in the total number of eyespots among adult individuals. The only source of variation in eyespot number would be the annual addition of eyespot feathers, in rows of 10 or 11, giving rise to variation between age classes.

These results and other observations on the museum samples led us to three insights that would help explain conflicting results between studies in the literature. First, eyespot number and train size are developmentally connected such that eyespots do not drive train length: it is the other way around—it is the feather/train growth that pulls eyespots up, spreads them out, and makes them visible to the female from afar. Second, we showed that eyespot number and male tail growth both have asymptotic functions, which means that later stage addition of feathers would lead to minor and ineffective eyespots with no added benefit to males. Finally, we argue that male vigor is a crucial factor modulating the effects of male size and beauty. Two males can have the same number of eyespots but differ in their age and vigor. Taken together, these insights can explain many of the conflicting results reported from different studies in the past.

Females prefer large ornaments (Summers and Ord, 2022) but that apparent “preference” can be due to male-driven, female-driven, or male- and female-driven causes. Based on our results and insights, we have proposed a two-stage, multimodal model of female choice based on male size, beauty, and vigor and suggest that it is not the number of eyespots but the size of the eyespot-studded-train that may be the basis of female choice. Females may be attracted by train size from afar and male quality and male vigor and display may come into play when they are in close proximity. We conclude that females may be choosing the tallest, most vigorous, and most “beautiful” males. Our results solve the problem of the relationship between eyespot number and tail length and provide a new perspective on the role of male size, vigor, and beauty in female choice, and a new avenue for mounting research on the development and evolution of the peacock’s tail.

## Acknowledgements

We thank Kevin Kerr (Toronto Zoo), Mark Peck (ROM), Paul Sweet, and Lydia Gaetano (AMNH) for their help with our work at the museum. We also extend our sincere thanks to Ellen Larsen, Daniel Hartl, Richard Morton, Bhagwati Gupta, Jonathan Stone, and Dave Rollo for their comments on the manuscript. We owe our gratitude to Robert Montgomerie and Roslyn Dakin for their expert, critical comments on the manuscript. We would like to thank all the anonymous reviewers of the earlier versions of this manuscript; their comments have made us think deeply and look for answers to explain the contradictory results reported from different studies. Michelle Brown provided technical support. This work was supported by grants from a Natural Sciences and Engineering Research Council of Canada and McMaster University to RSS.

## Author contributions

RSS: Development of concept, collection of data, theoretical–developmental modelling, critical review of sexual selection theories, preparation of the manuscript, and financial support; SJ: literature review, critical analysis of sexual selection theories, data analysis, and preparation of the manuscript.

## Data sharing plan

The original data obtained on the museum specimens can be obtained from: Singh and Jagadeeshan, 2021. Dryad My Dataset: doi: 10.5061/dryad.1g1jwstwg

